# Genome-wide analysis of ivermectin response by *Onchocerca volvulus* reveals that genetic drift and soft selective sweeps contribute to loss of drug sensitivity

**DOI:** 10.1101/094540

**Authors:** Stephen R. Doyle, Catherine Bourguinat, Hugues C. Nana-Djeunga, Jonas A. Kengne-Ouafo, Sébastien D.S. Pion, Jean Bopda, Joseph Kamgno, Samuel Wanji, Hua Che, Annette C. Kuesel, Martin Walker, Maria-Gloria Basáñez, Daniel A. Boakye, Mike Y. Osei-Atweneboana, Michel Boussinesq, Roger K. Prichard, Warwick N. Grant

**Affiliations:** Department of Animal, Plant and Soil Sciences, La Trobe University, Bundoora, 3086, Australia; Wellcome Trust Sanger Institute, Hinxton, Cambridge, CB10 1SA, United Kingdom; Institute of Parasitology, McGill University, 21111 Lakeshore Road, Sainte Anne-de-Bellevue, Québec, H9X3V9 Canada; Parasitology and Ecology Laboratory, Department of Animal Biology and Physiology, Faculty of Science, P.O. Box 812, University of Yaoundé 1, Yaoundé, Cameroon; Centre for Research on Filariasis and other Tropical Diseases (CRFilMT), P.O. Box 5797, Yaoundé, Cameroon; Research Foundation in Tropical Diseases and the Environment (REFOTDE), P.O. Box 474, Buea, Cameroon; Institut de Recherche pour le Développement (IRD), IRD UMI 233 TransVIHMI – Université Montpellier – INSERM U1175, 911 avenue Agropolis, P.O. Box 64501, 34394 Montpellier Cedex 5, France; Faculty of Medicine and Biomedical Sciences, University of Yaoundé 1, P.O. Box 1364, Yaoundé, Cameroon; UNICEF/UNDP/World Bank/World Health Organization Special Programme for Research and Training in Tropical Diseases (WHO/TDR), World Health Organization, 20, avenue Appia, 1211, Geneva, Switzerland; London Centre for Neglected Tropical Disease Research, Department of Infectious Disease Epidemiology, Faculty of Medicine, School of Public Health, Imperial College London, W2 1PG, United Kingdom; Noguchi Memorial Institute for Medical Research, University of Ghana, P.O. Box LG 581, Legon, Ghana; Council for Scientific and Industrial Research (CSIR), Department of Environmental Biology and Health Water Research Institute (CSIR), P.O. Box M 32, Accra, Ghana

**Author notes:** Corresponding authors: Stephen R Doyle, Roger K Prichard, Warwick N Grant. These authors contributed equally to this work (Stephen R. Doyle, Catherine Bourguinat).

**Keywords:** *Onchocerca volvulus*, ivermectin, Illumina genomics, Pool-seq, sub-optimal drug response, population genetics, genetic drift, soft selection

## Abstract

**Background:** Treatment of onchocerciasis using mass ivermectin administration has reduced morbidity and transmission throughout Africa and Central/South America. Mass drug administration is likely to exert selection pressure on parasites, and phenotypic and genetic changes in several *Onchocerca volvulus* populations from Cameroon and Ghana - exposed to more than a decade of regular ivermectin treatment - have raised concern that sub-optimal responses to ivermectin’s anti-fecundity effect are becoming more frequent and may spread.

**Methodology/Principal Findings:** Pooled next generation sequencing (Pool-seq) was used to characterise genetic diversity within and between 108 adult female worms differing in ivermectin treatment history and response. Genome-wide analyses revealed genetic variation that significantly differentiated good responder (GR) and sub-optimal responder (SOR) parasites. These variants were not randomly distributed but clustered in ~31 quantitative trait loci (QTLs), with little overlap in putative QTL position and gene content between countries. Published candidate ivermectin SOR genes were largely absent in these regions; QTLs differentiating GR and SOR worms were enriched for genes in molecular pathways associated with neurotransmission, development, and stress responses. Finally, single worm genotyping demonstrated that geographic isolation and genetic change over time (in the presence of drug exposure) had a significantly greater role in shaping genetic diversity than the evolution of SOR.

**Conclusions/Significance:** This study is one of the first genome-wide association analyses in a parasitic nematode, and provides insight into the genomics of ivermectin response and population structure of *O. volvulus*. We argue that ivermectin response is a polygenically-determined quantitative trait in which identical or related molecular pathways but not necessarily individual genes likely determine the extent of ivermectin response in different parasite populations. Furthermore, we propose that genetic drift rather than genetic selection of SOR is the underlying driver of population differentiation, which has significant implications for the emergence and potential spread of SOR within and between these parasite populations.

**Author summary:** Onchocerciasis is a human parasitic disease endemic across large areas of Sub-Saharan Africa, where more that 99% of the estimated 100 million people globally at-risk live. The microfilarial stage of *Onchocerca volvulus* causes pathologies ranging from mild itching to visual impairment and ultimately, irreversible blindness. Mass administration of ivermectin kills microfilariae and has an anti-fecundity effect on adult worms by temporarily inhibiting the development *in utero* and/or release into the skin of new microfilariae, thereby reducing morbidity and transmission. Phenotypic and genetic changes in some parasite populations that have undergone multiple ivermectin treatments in Cameroon and Ghana have raised concern that sub-optimal response to ivermectin’s anti-fecundity effect may increase in frequency, reducing the impact of ivermectin-based control measures. We used next generation sequencing of small pools of parasites to define genome-wide genetic differences between phenotypically characterised good and sub-optimal responder parasites from Cameroon and Ghana, and identified multiple genomic regions differentiating the response types. These regions were largely different between parasites from both countries but revealed common molecular pathways that might be involved in determining the extent of response to ivermectin’s anti-fecundity effect. These data reveal a more complex than previously described pattern of genetic diversity among *O. volvulus* populations that differ in their geography and response to ivermectin treatment.

## INTRODUCTION

*Onchocerca volvulus* is a filarial nematode pathogen responsible for causing human onchocerciasis. The infection is associated with significant morbidity, ranging from itching to severe dermatitis and from visual impairment to blindness. This morbidity and its economic impact have motivated large scale disease control programmes in the foci located in South and Central America, Yemen and throughout Sub-Saharan Africa, where more than 99% of the global at-risk population, estimated at 100 million people, live. Currently, onchocerciasis control is based primarily on annual community directed treatment with the macrocyclic lactone, ivermectin (CDTI).

Ivermectin has a least two pronounced effects on the parasites: (i) an acute “microfilaricidal effect” that results in the rapid and almost complete removal of microfilariae—the larval progeny of adult worms—from the skin within days to weeks after treatment, and (ii) a sustained “anti-fecundity effect” that results in prolonged but temporary inhibition of the release of new microfilariae from adult female worms into the skin for approximately three to six months [1]. Although some reports describe an increased proportion of adult dead worms following multiple ivermectin treatment rounds [2, 3], ivermectin is generally considered not to be a macrofilaricide [4, 5]. Ivermectin mass treatment therefore aims to: (i) prevent, and to the extent possible revert, pathology by removing microfilariae from the skin and eyes and delaying repopulation of these tissues with new microfilariae, and (ii) reduce transmission of *O. volvulus* by reducing the number of microfilariae that can be ingested by the blackfly (*Simulium*) vectors. Biannual (6-monthly) mass administration of ivermectin in hypo- and meso-endemic areas, and three-monthly ivermectin administration in hyperendemic areas of Central and South America have or are likely to have permanently eliminated transmission in most foci of the Americas [6–12]. Annual CDTI (expanded to biannual CDTI in some cases) and/or vector control, have or are likely to have also eliminated onchocerciasis in a number of endemic areas in Africa [13–19]. The World Health Organization Roadmap on Neglected Tropical Diseases has set the ambitious target of achieving onchocerciasis elimination where feasible in Africa by 2020 [20], and the African Programme for Onchocerciasis Control proposed expanding this goal to 80% of the countries by 2025 [21].

In some areas of Africa, however, persistent microfilardermia (microfilaria in the skin) and transmission have been reported after 15–20 years of ivermectin treatment [22–33]. Already in 2004, an *O. volvulus* ivermectin response phenotype termed “sub-optimal response” (SOR), characterized by the presence of live stretched microfilariae in the uteri of the adult worms 90 days after treatment, and consequent repopulation of the skin with microfilariae earlier/more extensively than expected based on prior data, was described in Ghana [29, 30]. Additional investigations have reported this phenotype also in other areas in Ghana [27, 34, 35] and in Cameroon [36, 37]. Allele frequency change in a number of “candidate” ivermectin response genes (chosen for analysis based on specific hypotheses concerning mechanisms of resistance to the acute effects of ivermectin in *O. volvulus*) has also been demonstrated in these populations when sampled before and after several rounds of ivemectin treatment [38–44], which is consistent with population genetic changes associated with drug selection pressure. While genetic selection for SOR was not demonstrated, these studies suggest that these populations are being influenced at the genetic level by ivermectin treatment and, if these “candidate” genes mediate the phenotypic changes in ivermectin response, that SOR has a genetic basis that may involve selection on several genes.

The reproduction biology and transmission dynamics of the parasite after ivermectin treatment have been described [5, 36, 37, 45]. However, further work is required to understand: (i) the variability in response to ivermectin that has been observed in ivermectin naïve parasite populations [46–48], (ii) the biological mechanism(s) by which parasites may tolerate and/or actively mitigate the inhibitory effects of ivermectin, and (iii) the corresponding genetic architecture underpinning these biological mechanisms, the influence of genetic selection, and the potential for SOR genotypes to be transmitted preferentially within and between parasite populations and thus increase in frequency. Although there has been some debate regarding the existence of SOR to ivermectin in *O. volvulus* [49–53], modeling of SOR – using individual-patient data on the rate of skin repopulation by microfilariae following treatment in communities with different histories of control [27, 54] – has provided support for the conclusion that the early reappearance of microfilariae in the skin that defines SOR is most likely due to a decreased susceptibility of the parasite to ivermectin’s anti-fecundity effect. Onchocerciasis morbidity reduction is driven primarily by ivermectin’s microfilaricidal effect but its anti-fecundity effect, delaying repopulation of the skin with microfilariae, also contributes. More importantly, since both of these effects are critical for achieving interruption of parasite transmission, an increase in the frequency of SORs could jeopardise onchocerciasis elimination goals [55].

Genome-wide approaches are increasingly being employed to investigate the effects of drug selection in human pathogens, including but not limited to, *Plasmodium falciparum* [56–60], *Schistosoma mansoni* [61], *Leishmania donovani* [62] and *Salmonella enterica* Typhi [63]. Such approaches have been instrumental in both confirming known and identifying novel drug resistance-conferring loci in experimental and natural populations, and in clarifying the roles played by selection, parasite transmission and genetic drift in drug responses. An important feature of these studies is that they do not rely on assumptions concerning mechanisms of resistance or candidate resistance genes. The unbiased whole-genome approach has proven particularly useful where there is a plausible hypothesis of polygenic inheritance of a quantitative trait (QT, i.e. a trait that is determined by interactions between multiple genes and the environment, and therefore will have a continuous distribution of phenotypic values in a population) and when the analysis is based on natural, outbreeding field populations in a non-model species in which the population genetic structure is unknown but may confound simple candidate gene analysis. Given that a number of different candidate genes have been proposed to be associated with genetic response to ivermectin treatment in *O. volvulus* and other parasitic nematode species (see reviews from Gilleard [64] and Kotze *et al*. [65] for discussion of success and limitations of candidate gene approaches), the hypothesis that variation in response by the parasite to ivermectin is a polygenic quantitative trait is both plausible and likely. Analyses to understand SOR have been limited by the fact that *O. volvulus* is genetically diverse [66], populations are likely to be genetically structured at an unknown spatial scale [67, 68], and the parasite is a non-model outbreeding organism that is not amenable to more direct controlled genetic crosses and quantitative trait loci (QTL) mapping methods. These challenges are further exacerbated by limitations in providing an accurate estimate of the degree of SOR in a given individual or population; microfilarial density in the skin determined by skin snip is an indirect estimate of SOR and is not precise, especially when the parasite density in the skin is low due to ivermectin treatment, and analysis of the reproductive status of adult worms by embryogram is typically (but not exclusively) qualitatively measured although it is a quantitative trait. However, recent advances in our understanding of the genome [69] and transcriptome/proteome of *O. volvulus* [70], and more broadly, investment in publicly available helminth biology resources such as WormBase Parasite (http://parasite.wormbase.org/, [71]), have provided the means by which a comprehensive evaluation of ivermectin-mediated selection of drug response in *O. volvulus* may now take place.

In this study, we present a genome-wide genetic analysis of drug response by comparing genetic diversity within and between pools of adult *O. volvulus* from Cameroon and Ghana that have been classified as “ivermectin-naïve or little treated” [NLT]), “good responder” (GR), and ivermectin SOR based on host population and/or individual host treatment history, microfilarial repopulation in the host skin after ivermectin treatment and embryogram analysis of female worms. This analysis has provided new insight into the putative genetic and biological mechanism(s) of response by *O. volvulus* to ivermectin. Underlying population structure, low susceptibility to ivermectin’s anti-fecundity effect, and potential for increase in the frequency of such phenotypes are discussed in the context of efforts to control and eliminate onchocerciasis in Africa.

## MATERIALS & METHODS

### Sample history and phenotyping of ivermectin response

*O. volvulus* samples used in this study were acquired from previously described studies conducted in Cameroon [2, 36, 37] and Ghana [34, 35]. Ethical clearances were obtained for the original sampling of parasites from the National Ethics Committee of Cameroon (041/CNE/MP/06), the Cameroon Ministry of Public Health (D30-65/NHA/MINSANTE/SG/DROS/CRC and D31/L/MSP/SG/DMPR/DAMPR/SDE/SLE), the Ethics Committee of Noguchi Memorial Institute for Medical Research, (NMIMR-IRB CPN 006/01-04) and McGill University (FWA 00004545).

Two separate experiments utilizing adult female worms collected from people with known individual and/or community ivermectin treatment history and response phenotype are reported here (see **Additional file 2: Figure S1** for maps of sampling regions in each country). Phenotypic classification of ivermectin response has been described in detail previously [34–37], and for this study was determined based on a combination of host level (skin microfilariae density) and female worm level (presence or absence of stretched microfilariae *in utero*) characteristics (**Additional file 2: Table S1**).

A total of 108 adult female parasites were used in the Pool-seq analyses; a description of the samples used, phenotypic characterization, host and host-community ivermectin treatment history is presented in **Additional file 2: Table S2**. Briefly, Pool-seq samples from Cameroon consisted of 3 pools of parasites composed of: (i) ~40 “drug-naïve or little treated” (NLT) female worms from the Nkam valley, (ii) 22 ivermectin GR worms, and (iii) 16 SOR worms. The Cameroon GR and SOR parasites were collected from people living in the Mbam valley who live in communities that have participated in mass drug administration of ivermectin since 1994 (at least 13 years prior to sampling). In addition to these treatments, these people from which parasites were collected also received between 4 to 13 doses of ivermectin between 1994 and 1997 in a controlled clinical trial [2]. Similarly, the Pool-seq samples from Ghana consisted of 3 pools of parasites made up from: (i) 10 worms categorized as NLT, having been exposed to ivermectin 455 and 90 days prior to the time of sampling, (ii) 7 GR worms exposed to 11–16 annual doses of ivermectin prior to sampling, and (iii) 13 SOR worms who had been exposed to 9–16 annual doses of ivermectin.

A total of 592 adult female worms were used in the single worm genotyping experiments; a summary of the worms selected for Sequenom genotyping as well as the treatment history of the hosts and the area in which the hosts live are described in **Additional file 2: Table S3**. Parasites from Cameroon (n = 436) are categorized in three groups of samples: (i) the Nkam Valley (NKA07; n = 140), from individuals who had been exposed to only a single ivermectin treatment 80 days prior to sampling, i.e., considered NLT [37], (ii) Mbam Valley (MBM07; n = 112), from people who had been exposed to multiple rounds of ivermectin treatment prior to sampling as described above [37], and (iii) parasites from individuals in the Mbam Valley who were truly ivermectin naïve (MBM94; n = 184), sampled in 1994 prior to introduction of CDTI [2]. Response phenotype data are not available for the MBM94 parasites. In total, 184 ivermectin naïve with unknown phenotype, 225 GR, and 27 SOR parasites were genotyped. Worms from Ghana (n= 156) used for Sequenom genotyping were sampled from 6 communities with different ivermectin exposure histories (range: 2–17 treatments; **Additional file 2: Table S3**), and were composed of 105 GR and 43 SOR parasites. An additional 8 parasites that had been exposed to multiple annual ivermectin treatments but whose phenotype is not available were also included.

### DNA preparation and genome resequencing

DNA extraction from worms from Cameroon was performed in the REFOTDE laboratory in Cameroon using an EZNA tissue DNA kit (Omega Bio-Tek Inc., Norcross, GA, USA). DNA extraction from worms from Ghana was performed using the DNeasy^®^ Blood and Tissue kit (Qiagen®, Toronto, ON, Canada) at McGill University in Montreal, Canada.

The sequence data were generated from pools of worm DNA that were prepared by combining the DNA of individual worms that shared drug phenotype and geographic origin (i.e. NLT, GR or SOR from Ghana in 3 pools; NLT, GR and SOR from Cameroon in 3 pools; **Additional file 1: Table S1**). DNA was pooled from individual worms at McGill University and sequencing was carried out at the McGill University and Génome Québec Innovation Centre (Montreal, Canada) across 8 Illumina GAII lanes (Cameroon: GR and Ghana: SOR pools had sufficient DNA to allow two sequencing lanes to increase sequencing depth). Overall, ~270-million 76-bp single-end sequences were generated, resulting in an estimated 35.18-fold (standard deviation [SD] ± 13.30) unmapped coverage per pool (or ~0.65–4 fold per worm if equal amounts of DNA/worm are assumed). Sequence data are archived at the European Nucleotide Archive (ENA) under the study accession PRJEB17785.

### Read mapping and variant calling

Reads from each pool were aligned to the draft genome sequence O_volvulus_Cameroon_v3 (WormBase Parasite; [69]) using BWA-MEM [72], after which reads were aligned around putative indels and duplicate reads removed using GATK (v3.3-0) [73]. Approximately 70% of the raw data were mapped, resulting in an average mapped coverage of 24.47 ± 9.07 per pool. Three ‘pooled-sequencing’ aware variant calling approaches were used to analyse the pooled mapping data: CRISP [74], FreeBayes (using the “pooled-continuous” parameter) (https://github.com/ekg/freebayes) and PoPoolation2 [75](**Additional file 1: Table S2**). To ensure that comparisons could be made between all sequencing pools, the raw sequence nucleotide polymorphism (SNP) data were filtered using the following criteria: (i) the mapped read depth across a variant site was at least 8 reads but not greater than 3 SD from the genome-wide mean read depth for the given pool; (ii) there was no evidence of significant strand bias between forward and reverse reads; (iii) variants were bi-allelic SNPs, and (iv) variants associated with reads mapped to *Wolbachia* or mtDNA sequence were removed. The intersection of these three approaches post-filtering yielded 248,102 shared variable sites relative to the reference sequence.

### Genome-wide analyses of differentiation

To perform the genome-wide analyses of genetic differentiation of the sequencing pools, we used only variants found in the intersection of the three Pool-seq-aware SNP calling approaches described above for all following analyses. Given the statistical uncertainty associated with interpreting low coverage read frequencies as a proxy for allele frequency for any given SNP locus, and in turn, the need to use heavily-filtered variant call sets to enable comparison between groups, we focused on analyses of relative genetic diversity between groups by calculating F_ST_ in 10-kb sliding windows across the genome, and F_ST_ of gene features based on the O_volvulus_Cameroon_v3 genome build from WormBase Parasite (http://parasite.wormbase.org/; onchocerca_volvulus.PRJEB513.WBPS5.annotations). Both datasets were generated using PoPoolation2 with the following parameters: --min-count 2, --min-coverage 8, --max-coverage 2%, with corresponding individual haploid pool sizes specified). Analyses of significance between different groups for individual SNPs were performed using Fisher’s exact tests to explore shared variants present in both Ghana and Cameroon. Analyses of variant frequency were performed using the variant read frequency generated from the CRISP output.

One of the primary aims of this analysis was to define the physical map positions of regions of the genome that differed significantly between phenotypic classes in each country, on the assumption that such a region would contain a locus or loci that contributed to the phenotypic difference between the pools in question. We defined a genomic location as significant if it consisted of two or more adjacent 10-kb windows (from the F_ST_ sliding window analysis) in which the GR *vs* SOR F_ST_ values were greater than 3 SDs from the genome-wide mean Fst within a 50-kb window (unless otherwise stated).

### Single worm genotyping by Sequenom® MassARRAY

A subset of genome-wide SNPs (160 in total) were chosen to explore ivermectin association and population structure further by Sequenom genotyping of individual adult female worms phenotyped for ivermectin response. DNA from 592 individual female worms (described in **Additional file 2: Table S3**) was prepared for Sequenom® MassARRAY genotyping (Sequenom, Inc., San Diego, CA, USA)[76]. Due to the DNA quantity requirements for Sequenom analysis (600 ng per sample), many individual worm DNA samples (401 of 436 samples from Cameroon, 96 of 156 samples from Ghana) required whole genome amplification to increase the DNA concentration. This was performed using the REPLI-g screening kit (Qiagen®,Toronto, ON, Canada). DNA concentrations of all samples were quantified using Quant-iT™ Pico Green dsDNA Assay Kit (Life Technologies Inc, ON, Canada), before sending to the McGill University and Génome Québec Innovation Centre for genotyping.

### Population- and ivermectin association analyses

Sequenom data were analysed using PLINK [77]. Raw data were filtered based on allele frequency (loci with <0.05 minor allele frequency were removed; PLINK --success rate were removed3maf 0.05), genotype quality (samples with <80% assay success rate were removed: PLINK --mind 0.2; SNPs with <80% genotype call frequency were removed: PLINK --geno 0.2). Filtered Sequenom data (130 SNPs in 446 samples [**Additional file 2: Table S3**; sample numbers in analysis are indicated in parentheses]; 81.25% of total SNPs and 75.34% of total samples, respectively) were analysed by multidimensional scaling to assess geographic vs phenotypic influence on genetic diversity. Hardy-Weinberg equilibrium (HWE) was analysed using the same filtering conditions described above (PLINK --hardy), using the “--keep” function to analyse samples from each country separately. Discriminant analysis of principal components (DAPC) and population assignment based on membership probabilities were performed using the R package *adegenet* [78].

## RESULTS and DISCUSSION

### Genome resequencing of pooled, phenotyped O. volvulus from Ghana and Cameroon

A genome-wide approach was used to detect genetic signatures associated with the “sub-optimal responder” (SOR) phenotype of *O. volvulus* when exposed to ivermectin. The criteria used to differentiate the SOR phenotype from the “good responder” (GR) phenotype were based on criteria described previously [29, 30]. In the present study, GR and SOR differ in two (deemed to be related) ways: (i) at the host level, the number of microfilariae in the skin determined by diagnostic skin snips is >7% of the pre-treatment value in SOR but microfilariae are largely undetectable in GR around 3 months post treatment (80 or 90 days for the samples from Cameroon and Ghana, respectively), and (ii) at the parasite level, SOR macrofilariae contain “stretched” microfilariae (ready to be released) *in utero* around 3 months post treatment (80 or 90 days for the samples from Cameroon and Ghana, respectively), as determined by embryogram, while GR do not. A SOR parasite, therefore, causes earlier repopulation of the skin with microfilariae than a GR parasite due to an earlier resumption of microfilarial release after temporary inhibition of fecundity.

To investigate the genetic differences between GR and SOR adult *O. volvulus*, whole genome sequencing was performed on pools of adult female worms from Cameroon and Ghana that were classified as “naïve or little treated” [NTL], or multiply treated GR and SOR groups based on the prior ivermectin exposure of the hosts and/or community from whom they were collected, and the host and parasite level response to ivermectin as described above (see **Additional file 2: Table S1** for response criteria and **Additional file 2: Table S2** for characteristics of the worm pools). We identified 248,102 variable positions that were shared among all groups and passed our stringent filtering criteria for inclusion in a genome-wide scan of genetic differentiation within and between pools (**Additional file 1: Table S2**), at an average marker density of 1 variant per ~389-bp (of the 96,457,494-bp nuclear genome). This represented only 32.7% of the total putative variants identified using the three variant analysis pipelines, primarily as a result of the relatively relaxed variant calling conditions of PoPoolation2 (**Additional file 1: Table S2;** 34.4% of SNPs called by PoPoolation2 were unique to this tool under the conditions used compared to 2.6% and 6.2% of unique SNPs called by CRISP and FreeBayes, respectively) and in part due to the stochastic variation in allele detection in low sequence coverage Pool-seq data.

Pool-seq has been used to estimate population genetic diversity in a number of different species [75, 79–82] on the assumption that individual read frequencies at a variant site are a proxy for allele frequency. This approach relies on sampling sufficient reads at any given position to be confident in detecting the diversity present; the more reads sequenced at a given position, the closer the read frequency is to the “true” allele frequency. Given that approximately 24.47-fold mapped coverage per pool (range: 0.53–3.05-fold coverage per worm) was obtained overall, it is unlikely that any of the pools was sequenced at sufficient depth to sample each genome present. Analyses of variation in read frequency between pools at low sequence coverage for any given nucleotide variant should, therefore, be treated with caution as they are confounded by significant statistical variation in coverage per genome [83]. In addition to strict filtering of the variants (e.g. to remove putative variant sites that were not detected by all three variant callers in all pools), we have attempted to account for this uncertainty by focusing on genetic variation calculated from multiple SNPs, either in sliding windows across the genome or on whole genes, rather than on individual nucleotide variants. A significant finding of this study was that genetic variation that differentiated GR and SOR pools was not randomly distributed but was strongly clustered in multiple discrete regions of genome. This observation establishes clearly, for the first time, both the genetic architecture and likely mode of selection of ivermectin response in *O. volvulus*, which is described in detail below.

### Genome-wide genetic differentiation between ivermectin response phenotypes

The two important questions with respect to the evolution of SOR are (i) which locus or loci are under selection, and (ii) whether the same loci are under selection in different populations, i.e., whether variation that differentiated SOR and GR worms in Ghana also differentiated SOR and GR worms in Cameroon. Shared variation between diverse geographic regions that differentiated SOR from GR would provide candidate markers that may predict ivermectin response in previously uncharacterised populations, and thus form the basis for the development of a ubiquitously applicable tool for monitoring the relative frequency of SOR and GR before and during CDTI. Pairwise analyses of individual SNPs (p-values from Fisher’s exact test; **Figure 1A**) and F_ST_ calculated from 10-kb windows (**Figure 1B**) both revealed a higher degree of differentiation between SOR and GR pools from Cameroon than between SOR and GR from Ghana, i.e., there were more loci or regions above significance thresholds in the Cameroonian pools. This difference is likely to be due to the unequal sample size between the two countries (Cameroon = 22 GR and 16 SOR worms; Ghana = 7 GR and 13 SOR worms); a greater proportion of total genetic diversity will be present in the Cameroon dataset as more worms are present, however, at the same time, the sequencing depth per Cameroon genome is lower than for the Ghana samples, and hence, will increase the stochastic variation in the Cameroon variant frequency. Although many single SNP loci from each country showed significant variation between SOR and GR pools when each country was analysed independently, only a single intergenic SNP (OM1b_7179218) was significantly different between pools and common to both countries (**Figure 1A**; red dot) after a Bonferroni genome-wide correction (**Figure 1A**; dashed lines) was applied. Furthermore, only six 10-kb regions that showed significant deviation above a genome-wide mean F_ST_ threshold of +3 SDs between GR and SOR were shared between Ghana and Cameroon (**Figure 1B**; dashed lines; red dots; **Table S3**). Relaxing the threshold to +2 SDs yielded 22 additional 10-kb regions that were able differentiate GR from SOR and were shared between countries (**Figure 1B**, orange dots; **Table S3**). A total of 28 F_ST_ windows at > + 2SDs is only marginally more than the number of windows that is expected to exceed +2 SDs by chance alone (0.05^2^ x 9893 10-kb loci = 24.73), and may in fact be inflated considering: (i) 11 of the 28 10-kb windows were immediately adjacent to at least one other 10-kb window in the genome and, therefore, are unlikely to all be segregating independently, and (ii) four of these 28 10-kb windows contained sequences associated with Pao retrotransposon peptidase-and integrase-related proteins and a further two windows contained ribosomal subunits (**Additional file 1: Table S3**). Given the multi-copy nature of these sequences, the high F_ST_ value for these 6 windows is likely to be a technical error associated with poor read mapping of multicopy sequences rather than true biological differentiation. These results suggest that, even at a reduced genome-wide level of significance threshold (i.e., > + 2 SDs), little variation that similarity discriminated GR from SOR parasites was shared between the two countries.

**Figure 1.**
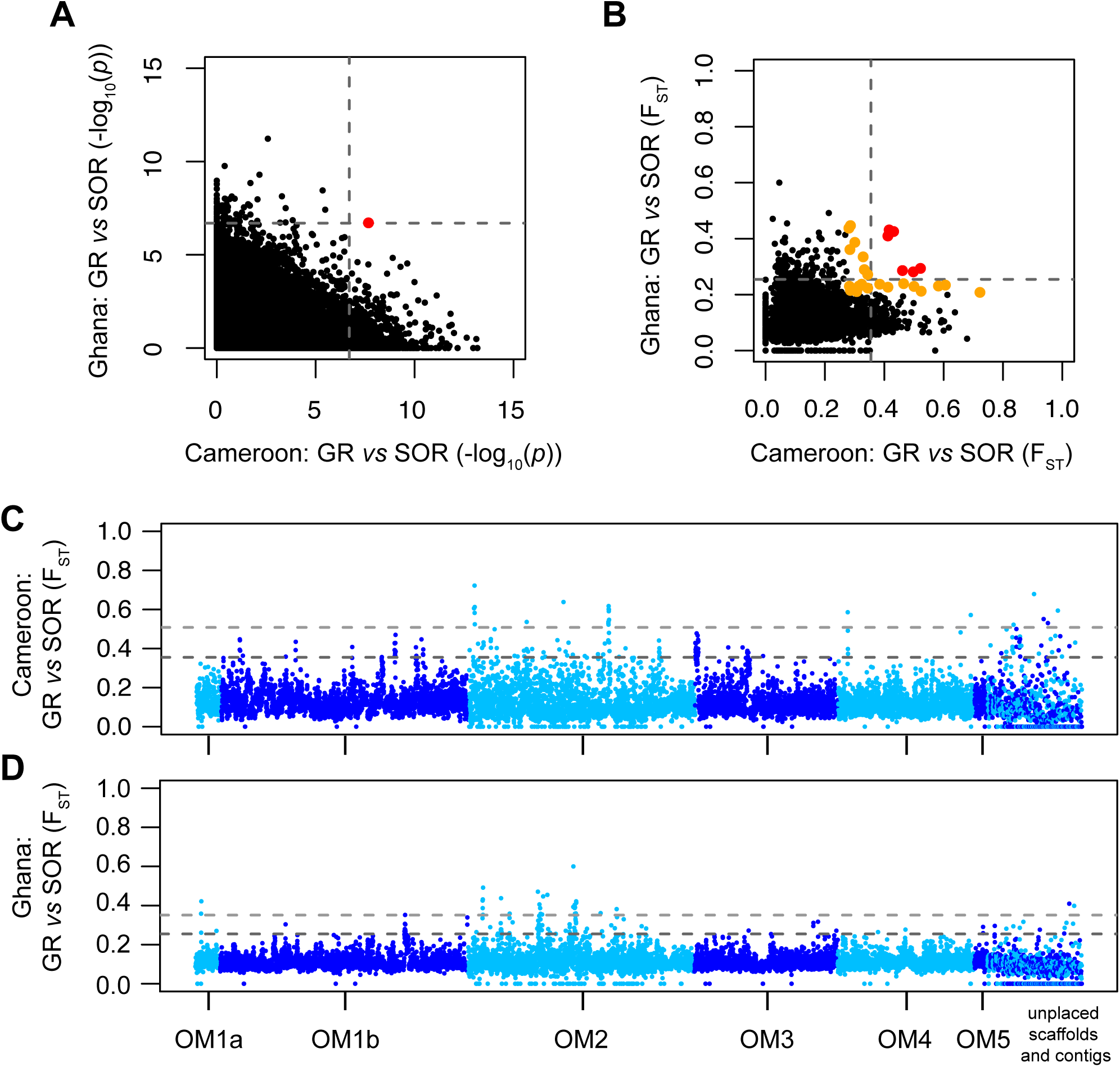
Analysis of shared variation that differentiates ivermectin good responder (GR) from sub-optimally responding (SOR) *Onchocerca volvulus* adult worms in both the Cameroon and Ghana population samples, measured using individual single nucleotide polymorphisms (SNPs) (Fisher’s exact test; **A**) or 10-kb windows (F_ST_; **B**). Dotted lines represent statistical cutoff applying the Bonferroni correction for SNPs and genome-wide mean F_ST_ + 3 standard deviations (SDs) (Cam: 0.355; Gha: 0.255) for 10-kb windows. Red dots highlight differentiation above genome-wide cutoffs that is shared by both groups. Orange dots represent additional shared differentiation at 2 SDs in the F_ST_ analysis (**B**). Manhattan plots of genome-wide F_ST_ describing spatial genetic differentiation between GR and SOR pools for both Cameroon (**C**) and Ghana (**D**). Each point represents F_ST_ calculated for a non-overlapping 10-kb window. Plots are colored to differentiate the main genomic scaffolds from unplaced scaffolds and contigs. Dotted lines represents genome-wide mean F_ST_ + 3 SDs (dark grey; Cam: 0.355; Gha: 0.255) and F_ST_ + 5 SDs (light grey; Cam: 0.508; Gha: 0.351).

To investigate the distribution of genome-wide genetic variation between ivermectin response groups, we analysed the relative positions of SOR vs GR 10 kb F_ST_ windows > +3SD in the *O. volvulus* genome. A striking finding was that the 10-kb regions that provided the most genetic differentiation between SOR and GR were not randomly distributed but were found in discrete clusters (**Figure 1C & D**); we interpret these clusters of significantly differentiated 10-kb windows (defined as 2 or more adjacent 10kb windows within a 50-kb window > +3 SDs from the genome-wide mean Fst calculated separately for each country) as putative QTLs. These QTLs are composed of a variable number of 10-kb windows and contain multiple genes. In total, 18 putative QTLs that mapped to well-assembled regions of the genome were found in the Cameroon data (**Figure 1C**; mean QTL size of 66,389-bp ± 55,157-bp SD) and 14 putative QTLs in the Ghana data (**Figure 1D**; mean size of 102,143-bp ± 95,690-bp SD), representing 1.23% and 1. 48% of the ~96.4-Mb nuclear genome, respectively (**Additional file 1: Table S4**). These data provide strong evidence that the SOR phenotype is a polygenic quantitative trait (QT) and, further, given that only a single putative QTL shared between Ghana and Cameroon was detected, suggest that different putative QTLs may be under selection in these two geographically separated parasite populations.

The apparent lack of concordance between Ghana and Cameroon SOR QTLs demonstrates that these two populations are genetically distinct. This observation, coupled with the number and location of putative QTLs, suggests that soft selection on pre-treatment genetic variation in the two parasite populations has acted on different loci in each country, and in turn, resulted in a different signature of selection in each country. Soft selection also implies that it may therefore be difficult to separate differences between Ghana and Cameroon SOR populations that are the result of selection from those that are the result of drift. However, the pattern of genetic variation within each country (see below) is consistent with the conclusion that the differentiation between GR and SOR worms (within the same country) is associated with phenotypic differences in ivermectin response (for background information on hard vs. soft selection, see **Additional File 2: Section 2**).

### Analysis of between-country genetic variation, and genetic diversity between responder phenotypes and “drug-naïve” worms

To test more explicitly for population structure between the two countries and thus, better understand the extent to which the standing genetic variation (see **Additional File 2: Section 2**) in these populations may have been shaped by the combination of selection and genetic drift, we analysed genetic differentiation between and within the two countries across all three treatment history/response categories (i.e. NLT, GR and SOR). A comparison of genome-wide allele frequency correlation within and between countries demonstrated that there was a significantly higher degree of shared variation (i.e., less population structure) within each country than there was between countries, where allele frequency correlation was low **(Figure 2A).** This supports the conclusion that there is significant genetic differentiation between the parasite populations in the two countries and that any genetic signal associated with SOR that might have been common to both countries would likely be masked by the presence of significant population genetic structure. Somewhat unexpectedly, a comparison of allele frequency correlations between response groups within each country suggested that, for both Cameroon and Ghana, the NLT worms were genetically more similar to the SOR worms than to the GR worms **(Figure 2A).** A direct pairwise comparison of the three treatment history/response categories within each country using genome-averaged F_ST_ values (calculated from 9,893 10-kb windows) was consistent with this observation: NLT *vs* SOR F_ST_ genomic medians were significantly smaller (Cameroon: 0.059; Ghana: 0.068) than either NLT *vs* GR medians (Cameroon: median = 0.095; two-sample Kolmogorov-Smirnov [KS] test: D = 0.0438, p < 0.001; Ghana: median = 0.110; KS D = 0.575, p < 0.001), or GR *vs* SOR medians (Cameroon: median = 0.112, KS D = 0.544, p < 0.001; Ghana: median = 0.104, KS D = 0.495, p < 0.001) (**Figure 2B**). The closer relationship between NLT and SOR than between NLT and GR is particularly surprising for Cameroon when one considers that the NLT and GR/SOR populations are from geographically distinct areas, i.e., the sampling sites within Nkam and Mbam Valleys were >100-km apart in two separate river basins separated by the Western High Plateau of Cameroon (see **Additional File 2: Figure S1 for map of sampling sites)**. However, seasonal dispersal of the local vector species, *Simulium squamosum*, has been observed in Cameroon over greater distances than the distance between the two sampling regions here [84], and therefore, some seasonal transmission between the two river basins may occur and requires further investigation.

**Figure 2.**
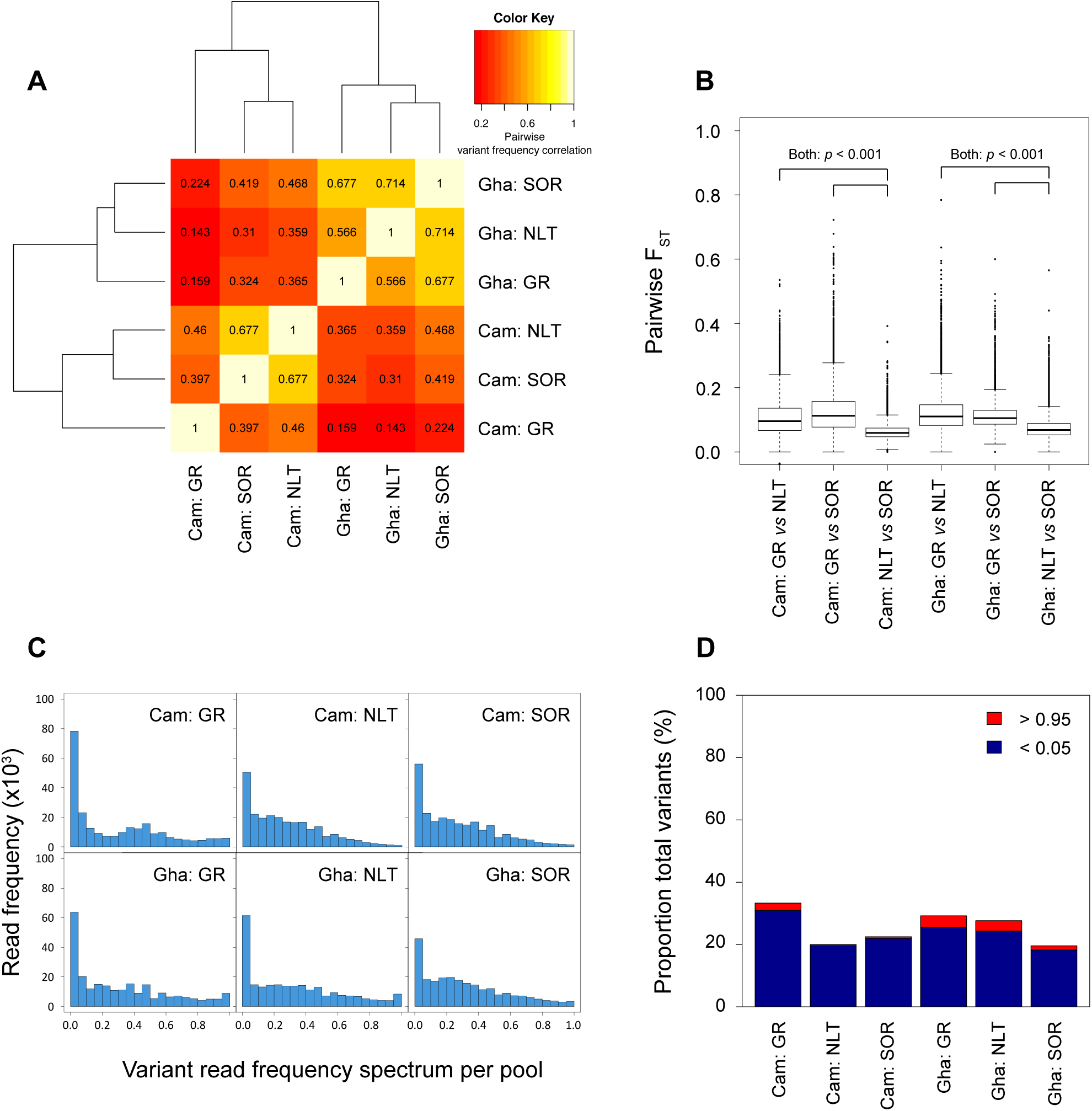
Analysis of genetic diversity between ivermectin responder phenotypes and drug-“naïve” (NTL) worms. (**A**) Spearman rank correlation analysis of variant read frequencies from 248,102 SNPs. Values within squares represents correlation coefficients for each pairwise analysis. (**B**) Degree of shared variation determined by pairwise comparisons of F_ST_ between treatment and response groups, summarized from 9893 10-kb windows throughout the genome. F_ST_ distributions were compared using a two-sample Kolmogorov-Smirnov [KS] test. (**C**) Variant read frequency spectrum from treatment and response subgroups for Cameroon and Ghana. The variant read frequency was calculated at each of the 248,102 SNP positions, from which the proportion of total variants in 0.05 frequency bins is presented. (**D**) Analysis of “invariant” loci per group as a proportion of the total number of variants observed, defined as variant read frequencies < 0.05 (blue) and >0.95 (red).

The observation of higher genetic similarity between the NLT and SOR populations supports the hypothesis of soft selection for SOR because it suggests that selection for SOR from an ivermectin naive population led to relatively little genome wide reduction in genetic variation. A characteristic prediction of soft as opposed to hard selection (see **Additional file 2: Section 2** for an extended discussion of soft-versus hard-selection) is that unlike hard selection, soft selection should not markedly reduce the genome-wide genetic diversity following selection. Since the SOR phenotype is associated with an early resumption of reproduction and, consequently, with the early availability of SOR offspring in the skin for vectors to ingest, ivermectin treatment likely interrupts the transmission of SOR genotypes for only a relatively short period of time between CDTI rounds compared to GR transmission interruption. If SOR alleles are already present on many different genetic backgrounds in the NLT starting population, this continued transmission of SOR genotypes under drug treatment will maintain genetic diversity in the SOR population, which is consistent with what we have observed in the present study.

The strong, genome-wide, between country genetic differentiation in the NLT populations implies that the standing genetic variation from which SOR is selected varies significantly between Ghana and Cameroon, and explains why the genetic outcome of selection of SOR differs between countries (as shown by the lack of concordance between Ghana SOR and Cameroon SOR populations). The presence of strong population structure between Cameroon and Ghana is not unexpected. The *O. volvulus* lifecycle is characterised by significant population bottlenecks at each transmission event: only a miniscule proportion of microfilariae in the skin are ingested by blackflies, and very few are subsequently transmitted to humans and establish as adult worms [85]. Repeated bottlenecks increase the severity of genetic drift by strongly enhancing the stochastic processes that generate genetic diversity between populations, independent of drug treatment, suggesting that genetic drift had created genetic differentiation between different parasite populations before initiation of CDTI, and that subsequent soft selection of SOR genotypes from these genetically distinct populations has led to SOR populations that are genetically distinct despite their phenotypic similarity.

In contrast, CDTI interrupts transmission of GR genotypes for a longer period of time. If this reduces the proportion of GR worms that contributes to the next generation, it will lead to a loss of genetic diversity and increase in genetic drift in subsequent GR populations relative to both the population before CDTI was initiated and the SOR population. This expectation is consistent with genetic change described in single-gene studies following treatment over time in *O. volvulus* [38–44].

To investigate the impact of soft selection on GR and SOR genetic diversity further, we estimated the genetic diversity within the group of GR worms and the group of SOR worms from each country in two complementary ways: (i) by calculating the variant read spectrum per group (as a proxy for the allele frequency spectrum, **Figure 2C**), and (ii) from the relative proportion of “invariant” SNP loci, which we have defined as SNP loci with variant read frequencies <0.05 or > 0.95 (**Figure 2D**). These complementary analyses provide insight into the degree of genetic variation present within each drug response parasite pool, and will detect loss of genetic diversity. The GR worms from both countries were less diverse and had a greater proportion of invariant loci than the SOR and NLT parasites, particularly in Cameroon, whereas the SOR and NLT parasites had similar levels of genetic diversity.

In conclusion, comparisons of genetic diversity and genetic similarity between NLT, GR and SOR support our hypothesis of soft selection for SOR, and explain both the relatively subtle signature of selection in the SOR parasites and the strong population structure observed between the SOR parasites from Cameroon and Ghana.

### Characterisation of molecular pathways identified from genes within QTLs that differentiate ivermectin response

In light of evidence suggesting that genetically separated parasite populations contain different standing genetic variation (likely to be the result of genetic drift) before introduction of CDTI, and thus that ivermectin-mediated soft selection may produce different genetic outcomes in SOR parasites between Cameroon and Ghana, it was of interest to compare genes within the putative QTLs identified in the SOR worms from Ghana and Cameroon. While we identified only a single putative QTL that was common to the SOR parasites from the two countries, the putative QTLs from each country contained genes encoding proteins that act in a limited number of molecular pathways. This implies that although different genes may be under selection in Ghana and Cameroon, there may be a common biological mechanism that confers SOR in both countries (see **Table 1** for a summary of genes with common functional characteristics and/or shared pathways identified in the QTLs, and **Additional file 1: Table S5** for characterization of all genes within each QTL).

**Table 1:** Summary of genes with shared functional characteristics / pathways from QTL peaks from Ghana and Cameroon ^a,b^

| Country | Ivermectin-associated | Neuro-transmission <sup>c</sup> | LIN-12/Notch signaling | Muscle assembly / myosin organization | Lipid synthesis and storage / stress |
| --- | --- | --- | --- | --- | --- |
| Cameroon |  | *Ovo-emc-6* (4; ACh)<br>*Ovo-cha-1* (9; ACh)<br>*Ovo-unc-17* (9; ACh)<br>*Ovo-stg-1* (10.1)<br>*Ovo-aex-3* (10.1; ACh)<br>*Ovo-nrfl-1* (16; ACh)<br>*Ovo-ncx-5* (24)<br>*Ovo-sca-1* (25)<br>*Ovo-lgc-46* (27; ACh) | *Ovo-crb-1* (4) | *Ovo-mup-4* (25)<br>*Ovo-mua-3* (25)<br>*Ovo-sca-1* (25)<br>*Ovo-mlc-5* (26) | *Ovo-acs-16* (3)<br>*Ovo-obr-2* (4)<br>*Ovo-math-46* (7)<br>*Ovo-hsp-17* (22)<br>*Ovo-mtd-15* (25)<br>*Ovo-cuc-1* (25) |
| Ghana | *Ovo-inx-5<sup>b</sup>* (5)<br>*Ovo-klp-11<sup>b</sup>* (13)<br>*Ovo-unc-44* (14) | *Ovo-nud-1* (1)<br>*Ovo-inx-5* (5)<br>*Ovo-stg-1* (10.2)<br>*Ovo-aex-3* (10.2; ACh)<br>*Ovo-unc-31* (13)<br>*Ovo-lgc-47* (17; ACh)<br>*Ovo-kin-2* (17)<br>*Ovo-unc-26* (19)<br>*Ovo-acc-1* (21; ACh)<br>*Ovo-snb-1* (23; ACh) | *Ovo-bre-5* (5)<br>*Ovo-pen-2* (29) | *Ovo-unc-82* (21)<br>*Ovo-nmy-1* (21) | *Ovo-hsb-1* (1)<br>*Ovo-sms-1* (13)<br>*Ovo-spl-1* (17)<br>*Ovo-fat-4* (19)<br>*Ovo-tat-2* (19) |

<sup>a</sup> Quantitative trait loci (QTL) identification is provided in parentheses (refer to **Additional file 1: Table S4** for a description of each QTL and **Additional file 1:** **Table S5** for the genes within). <sup>b</sup> putative ivermectin association as described in text <sup>c</sup> loci associated with acetylcholine neurotransmission are abbreviated with ACh.

The most prominent group of genes under the putative QTLs defined above were associated with neurotransmission (17 genes in 14 QTLs), which was encouraging considering that the primary target of ivermectin is a ligand-gated channel at neuromuscular junctions [86]. Given that the duration of the anti-fecundity effect of ivermectin distinguishes the GR and SOR phenotypes, the fact that nine of those genes (in eight QTLs) were associated with acetylcholine signaling is of particular interest because acetylcholine signaling plays an important role in the regulation of egg laying in *C. elegans*, and therefore, may be relevant to ivermectin’s anti-fecundity effect in *O. volvulus*. These genes include a number of ion channels (*Ovo-acc-1, Ovo-lgc-46, Ovo-lgc-47*), and components of acetylcholine synthesis (*Ovo-cha-1*), transport (*Ovo-unc-17, Ovo-aex-3*), and regulation (*Ovo-pha-2, Ovo-snb-1, Ovo-emc-6, Ovo-nrfl-1*). The *unc-17* and *cha-1* mutants in *C. elegans* exhibit hyperactive egg-laying phenotypes associated with defects in laying inhibition [87, 88], and acetylcholine activation of egg laying in *C. elegans* is regulated by neuropeptides, serotonin and glutamate [89]. Furthermore, acetylcholine receptors have recently been proposed to be involved in the development of the nervous system during embryo- and spermatogenesis in the filarial parasite, *Brugia malayi* [90]. In addition, two recent studies demonstrated inhibition of L-AChR receptors in *C. elegans* by ivermectin [91] and antagonistic effects of abamectin on nicotinic acetylcholine receptors [92], implying that under some circumstances ivermectin may act directly on acetylcholine signaling. We therefore hypothesise that modification of neurotransmission in general, and acetylcholine signaling pathways in particular, may contribute to ivermectin SOR and reflects an overall adjustment in neuromuscular signaling that mitigates the effects of ivermectin. This variation may also contribute to the changes in fecundity that have been associated with the SOR phenotype [93] by changing the way in which neurotransmission might influence the release of microfilariae in an analogous fashion to neuronal control of egg laying in *C. elegans*. Given the enrichment of neuronal genes that may play a role in regulating reproduction, it is of interest to note that the single QTL that is shared between Cameroon and Ghana SOR parasites includes *Ovo-aex-3*, a neuronal protein and regulator of synaptic transmission. This gene may be of particular interest because the *C. elegans* orthologue plays a role in reproduction and also in regulation of pharyngeal pumping [80] (a phenotype that is also consistent with reduced sensitivity to ivermectin in *C. elegans*). In addition, this QTL also includes *Ovo-stg-1*, which may be of interest due to its putative chaperone-like role in regulating ionotropic glutamate receptor (iGluR) function and a hypothesised role in protecting neurons from excitotoxicity or inappropriate depolarisation in *C. elegans* [81].

Nine putative QTLs contained genes associated with stress responses, including heat-shock proteins (*Ovo-hsp-17, Ovo-hsb-1*), and genes required for the synthesis (*Ovo-acs-16, Ovo-fat-4, Ovo-spl-1, Ovo-tat-2*), regulation and storage (*Ovo-obr-2, Ovo-sms-1, Ovo-math-46, Ovo-mtd-15, Ovo-cuc-1*) of lipids. Variation in gene expression in lipid metabolism-encoding genes following ivermectin treatment in *C. elegans* has been described previously [94], which was interpreted as a metabolic adaptation to starvation due to ivermectin inhibition of pharyngeal pumping but may also be a more general indicator of organismal stress. Genes associated in general with stress responses, including lipid metabolism, are often reported in genome-wide analyses for loci under drug selection pressure [56].

Other genes under the putative QTLs include those acting in pathways involved in developmental processes, including muscle assembly and myosin organization (three QTLs contained 6 loci; *Ovo-mup-4, Ovo-mua-3, Ovo-mlc-5*, *Ovo-unc-82, Ovo-nmy-1, Ovo-sca-1*), and germline and larval development signals associated with notch signaling (three QTLs), specifically with the suppression (*Ovo-bre-5*) or cleavage (*Ovo-pen-2,Ovo-crb-1*) of the LIN-12 receptor. The relevance, if any, of these developmental genes to ivermectin response is not known.

This list of genes that fall under the QTLs leads to a hypothesis of SOR as tolerance to ivermectin’s anti-fecundity effect brought about by a “re-tuning” of neuronal function in combination with a stress response. We acknowledge the tentative nature of this hypothesis given: (i) the limitations in the resolving power of genetic-association analysis based on modest sequencing depth of pooled samples, and only a single comparison between GR and SOR parasites from each country, (ii) the inability to carry out more specific analysis of coding sequence variation imposed by limited genetic resolution, (iii) the large number of predicted genes that lack functional annotation in the *O. volvulus* genome (~67.4% of the coding sequences within the QTLs are unannotated or hypothetical), and (iv) the methods available for phenotype classification, which has limited sensitivity for the determination of the presence and extent of microfilariae density in the skin [95] and is qualitative with respect to the presence or absence of live stretched microfilariae *in utero*, which in turn decreases the power of genetic association. However, the enrichment of neurotransmission and stress response genes in the QTLs that differentiate GR and SOR parasites in two geographically independent populations is striking, and does provide support for our hypothesis of a polygenic quantitative trait characterised by earlier recovery from the acute effects of ivermectin on fecundity.

If correct, this hypothesis implies that the SOR adult worms remain sensitive to the acute effects of ivermectin on fecundity but recover more quickly than GR worms. Early recovery from the acute effect of ivermectin on fecundity does not require a mechanism that is specific to the mode of action of that acute effect, so a polygenic “retuning” or “buffering” of the neuronal regulation of reproduction that leads to earlier recovery of fertility is biologically plausible. It may also explain why our analyses have failed to validate “candidate genes” that would protect worms against ivermectin’s acute effects (see below).

### Analysis of “candidate” ivermectin-resistance genes

Given the extensive literature focused on “candidate” genes (i.e. genes chosen for analysis based on specific hypotheses concerning mechanisms of resistance to the acute effects of ivermectin in *O. volvulus*), and their apparent absence from the QTLs described here, it was important to investigate these genes, which included glutamate-gated chloride channels (*Ovo-avr-14, Ovo-glc-2, Ovo-avr-15, Ovo-glc-4*), p-glycoproteins (*Ovo-pgp-1, Ovo-pgp-10, Ovo-pgp-11, Ovo-plp-1*), ABC transporters (*Ovo-abcf-1, Ovo-abcf-2, Ovo-abcf-3*) and other candidates (*Ovo-ben-1* [beta tubulin], *Ovo-lgc-37, Ovo-mrp-7, Ovo-dyf-7*). *Ovo-abcf-1* was the only “candidate” gene found in a QTL. For the majority of the other genes, no significant SOR *vs* GR Fst differences were observed (**Additional file 1: Table S6).** *Ovo-abcf-1* (in QTL-5), *Ovo-abce-1, Ovo-dyf-7, and Ovo-pgp-10* did show moderate levels of differentiation, but none were statistically significant. Given that our data suggests that parasite populations are structured and that alleles associated with response may vary between populations, we cannot exclude that these loci contribute to variability in ivermectin susceptibility in other *O. volvulus* populations. However, considering the evidence of multiple putative QTLs and strong geographic population structure, it is possible that single “candidate” gene comparisons may have been confounded by population structure and the polygenic nature of the trait. We conclude that there was no evidence of significant genetic differentiation in these candidate genes between the SOR and GR populations compared here.

Three of the 31 putative QTLs did contain “neuronal function” genes that have been linked previously to ivermectin sensitivity in *C. elegans* (**Table 1; Additional file 1: Table S5**): *Ovo-unc-44*, a likely orthologue of a *C. elegans* gene that influences ivermectin sensitivity and is involved in neuronal development [96], *Ovo-inx-5*, which in *C. elegans* encodes an innexin gap junction protein that is associated with the pharyngeal motor neurons and is related to *unc-9* (a known ivermectin-resistance allele [97]), and *Ovo-klp-11*, a kinesin motor protein found in the cilia of chemosensory neurons in *C. elegans*, which has significant homology to, and is a likely binding partner of, *Ovo-osm-3*, alleles of which have been described to decrease sensitivity to ivermectin in *C. elegans* [96].

### Geographic and genetic distribution of ivermectin susceptibility

Given the importance of the distinction between population structure as a result of genetic drift and phenotypic differentiation as a result of selection for SOR, we sought to investigate ivermectin response genetics in a larger cohort of individual female worms (most having been characterized for their ivermectin response phenotypes via embryograms [34–37]; **Additional file 2: Table S3**) by genotyping individual worms at 160 SNP loci by Sequenom (**Additional file 1: Table S7**). The loci were chosen based on the original Pool-seq analyses to determine the degree of genetic association with response phenotype, and to characterize genetic structure within these populations. Multidimensional scaling analysis (MDS) was used to interrogate the Sequenom genotyping data from 446 female worms at 130 SNPs that passed filtering criteria. Three aspects of the data are highlighted (in different panels, **Figure 3**): (i) between country population structure, (ii) within country population structure, and (iii) differentiation between response phenotypes.

**Figure 3.**
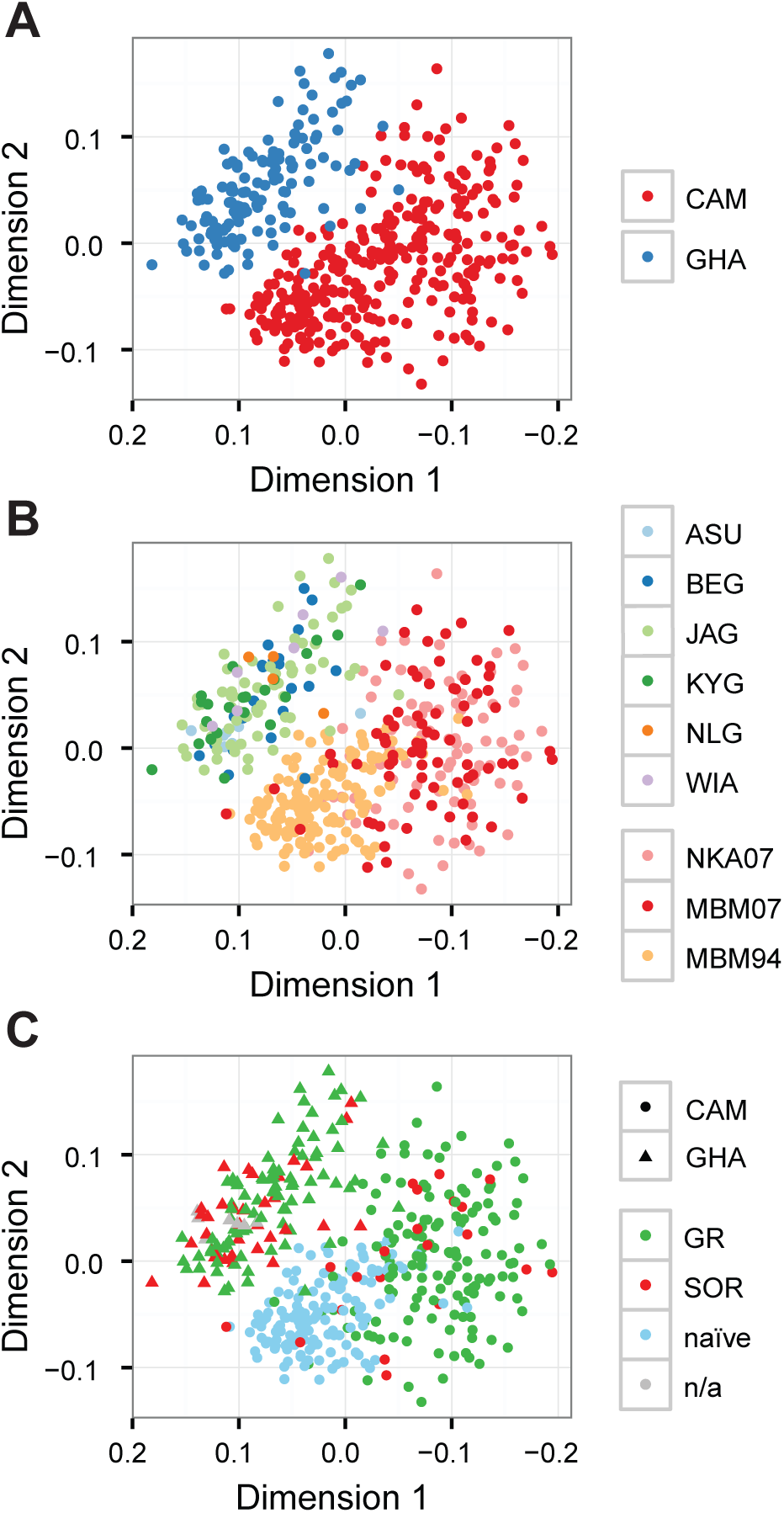
Analysis of genetic differentiation among 446 *O. volvulus* female worms from Ghana and Cameroon individually genotyped at 130 loci distributed throughout the genome. Multi-dimensional scaling analysis was used to determine the relative genetic similarity between worms. Plots contain the same data, but have been presented to emphasise the degree of genetic differences between countries (**A**), sampling communities within each country (**B**), and their treatment exposure and phenotypic response to ivermectin if known (**C**). Ghanaian sampling sites: Asubende (ASU), Begbomdo (BEG), Jagbenbendo (JAG), Kyingakrom (KYG), New Longoro (NLG), and Wiae (WIA). Cameroonian sampling sites: Nkam (NKA07), Mbam valley sampled in 1994 (MBM94) before introduction of annual CDTI in 1994 and sampled in 2007 (MBM07).

Consistent with the Pool-Seq analysis, we observed clear differentiation between worms from Ghana and worms from Cameroon: there were very few [blue] Ghanaian worms mixed with the [red] Cameroon worms (**Figure 3A**), regardless of response phenotype. Analysis of worms by their community of origin to allow within as well as between country comparisons did not reveal any internal genetic structure in the Ghanaian populations or between the Cameroonian NKA07 and MBM07 [pink and red, respectively] populations (**Figure 3B**).

Interpretation of the Cameroon data is more complex than for Ghana due to the likely existence of underlying population structure that is both temporal and spatial in origin: within Cameroon, MBM94 [light orange] was genetically distinct from both NKA07 and MBM07 populations but NKA07 and MBM07 were indistinguishable. MBM94 and MBM07 are two samples drawn from the same geographical location in Cameroon (the Mbam valley) at different times: MBM94 was sampled in 1994 before ivermectin distribution began, and is drug-naïve, whereas MBM07 was sampled in the Mbam Valley after 13 years of annual CDTI and contains approximately 16% SOR and 84% GR parasites (n = 112). Given that MBM07 is still largely GR, we interpret this signal of strong population differentiation as the result of genetic drift in MBM94, i.e., of largely stochastic (rather than deterministic, i.e., selection) genetic change likely brought about by multiple population bottlenecks imposed by the impact of 13 years of ivermectin treatment on parasite transmission and hence on effective population size. If this interpretation is correct, the striking similarity between MBM07 and NKA07 is coincidental: genetic drift is a stochastic process, and previously distinct populations may become more similar by chance alone.

Alternatively, the similarity between NKA07 and MBM07 may indicate that there is transmission between NKA07 and MBM07, such that as ivermectin shrank the originally naïve MBM94 population over 13 years, exchange of parasites between the Nkam Valley and the Mbam valley mediated by vector or human migration brought these two initially dissimilar populations closer together genetically. While the Nkam valley had not received CDTI at the time the NKA07 samples were collected (the samples genotyped are classified as NLT, having only been exposed to a single experimentally controlled round of ivermectin treatment [37]), some communities in the Nkam valley are only around 10 km from communities across the Nkam river which have received annual CDTI for >10 years. Consequently, the NKA07 parasites may not be entirely “naïve”. However, transmission of multiply-exposed parasites from these villages into NKA07 patients is unlikely to account for the similarities between NKA07 and MBM07, as the NKA07 population was largely composed of GR parasites (94.3%, n = 140).

Our data do not allow discrimination between these two hypotheses. In either case, however, the Cameroon data underline how little is known about the determinants of *O. volvulus* population structure in general, and the genetic impact of population bottlenecks imposed by ivermectin-mediated temporary interruption and long-term reduction of transmission in *O. volvulus*. These data also illustrate the confounding effect of underlying population structure on our ability to distinguish between selection and drift as drivers of genetic differentiation between two population samples, regardless of the origin of that structure, whether due to a drug induced population bottleneck, natural drift over time or parasite immigration from a drug naïve into a drug treated population.

Analysis of the distribution of response phenotypes revealed no clear separation of GR (green) from SOR (red) in either Ghana or Cameroon samples (**Figure 3C**). **Additional File 2 Table S3** and careful comparison of **Figures 3B** and **3C** show that the Cameroon NKA07 parasite group in **Figure 3B** (pink dots) is composed almost entirely of GR worms (SOR frequency of approximately 5%, n = 140), whereas the MBM07 parasite group (**Figure 3B**, red dots) is composed of a mix of GR worms (**Figure 3C**, green dots) and SOR worms (**Figure 3C**, red dots; SOR frequency approximately 16% [n = 112]), but that the GR and SOR worms in both NKA07 and MBM07 are intermingled, i.e. the differentiation between (NKA07 + MBM07) and MBM94 in **Figure 3B** is unrelated to the distribution of drug phenotypes in **Figure 3C**. Thus, these data suggest that genetic differentiation between the Cameroon populations is not correlated with response phenotype.

To further characterise the differences between GR and SOR parasites within each country, we analysed genotype frequencies for Hardy-Weinberg equilibrium (HWE). A greater number of SNPs were significantly out of equilibrium in the GR populations compared to the SOR populations from both countries (significance threshold of p < 0.05; 111 *vs* 20 (of 131) SNPs in the Cameroon GR *vs* SOR parasites, **Additional file 1: Table S8**; and 80 *vs* 68 (of 121) SNPs in the Ghana GR *vs* SOR parasites, **Additional file 1: Table S9**). In this respect, the Sequenom genotype and the whole genome PoolSeq data are concordant: greater deviation from HWE in the GR populations is best explained by ivermectin treatment reducing effective population size and genetic diversity and increasing genetic drift in the GR worms, but not in the SOR worms or naïve populations.

Collectively, these Sequenom data suggest that spatial (between Ghana and Cameroon) and temporal (pre-and post 13 years of drug exposure within Cameroon) genetic differentiation were readily detectable by genotyping at these 130 SNP loci, but that this genotyping failed to detect the relatively weaker signal of genetic differentiation between GR and SOR phenotypes. This is illustrated most clearly by comparison of the Ghanaian and Cameroon SOR populations (**Figure 3C**), where the MDS coordinates of Ghanaian SOR individuals cluster with the Ghanaian GR individuals, and Cameroon SOR individuals cluster with Cameroon GR, evidence that SOR alleles exist in genetic backgrounds determined by their population of origin, and that the genetic signature of soft selection for SOR in different naïve populations is weak compared to the pre-existing population structure that is the product of genetic drift. This is further supported by an analysis of genotype-based assignment of individual parasites to their respective populations (**Additional file 2: Section 3**). Greater that 99.5% of individuals were correctly assigned to their country of origin by their genotype, and between 40–92.4% of individuals from Cameroon and 0–97% of individuals from Ghana to their respective communities or regions. In contrast, 0% of Cameroon worms and only 8.1% of Ghanaian worms were correctly assigned as SOR worms on the basis of their genotypes, i.e., there is strong correlation between Sequenom genotype and place of origin but little or no correlation between Sequenom genotype and ivermectin response phenotype. Furthermore, the MDS analysis of the three Cameroon groups of samples (**Figure 3C**) supports the view that not only do past histories of geographic or ecological separation result in genetic drift and hence differentiation between *O. volvulus* populations, but that reduction or interruption of transmission and reduction in population size by ivermectin, and/or migration from untreated into treated populations also drives drift and masks selection.

### Implications for control and elimination

The success of CDTI in the majority of areas in Africa where CDTI has been implemented with good coverage [17], suggests that the alleles responsible for SOR are not common across Africa and that CDTI can interrupt transmission in many populations, depending on epidemiological and programmatic factors such as the intensity of blackfly biting on humans and the frequency and achieved coverage of CDTI [28, 33, 98–100].

However, it is also clear that drug response by *O. volvulus* is not uniform in all ivermectin-naïve populations [5, 27, 37, 46, 47], and that in some naïve populations (such as those sampled here), parasites with sub-optimal response to ivermectin’s anti-fecundity effect are present prior to ivermectin treatment. The data presented here provide an explanation for the presence of SOR worms in ivermectin-naïve populations, and support for the view that sub-optimal response to ivermectin in those populations is a genetically determined trait that can increase in frequency as a result of selection. Selection for SOR has progressed to an extent in the populations sampled for analysis here that the genetic signal of that selection can be detected in the Pool-seq data. The genetic signal is, however, weak because it is based on soft selection acting on many genes that contribute to a quantitative trait. The implications of these conclusions for control and elimination hinge on three crucial questions.

The first question is whether, and under what circumstances, SOR can be detected genetically. The whole genome Pool-seq data demonstrate that it is possible to detect SOR genotypes in an analysis of 248,102 SNPs but our first attempt to validate SOR alleles in individual worms by reducing the number of SNP’s to a panel 130 SNPs in Sequenom genotyping failed. This failure is likely because soft selection for SOR has left a faint genetic signature in the SOR populations compared to the very strong, pre-existing population structure signals and likely ivermectin-induced genetic drift in treated populations. More careful marker selection based on better quality whole genome sequence data from individual-rather than pooled-worms may solve this problem and permit sensitive detection of SOR genotypes in samples drawn from a single parasite population or transmission zone.

The second crucial question is whether it is possible to predict the likelihood that SOR frequency will increase to a level that prevents elimination of *O. volvulus* transmission in a given population. Our data suggest that ivermectin reduces the population size of GR worms, as one would expect, but that the population of SOR worms is stable and/or increasing as the GR population shrinks. In very simple terms it is, therefore, a race: will the rate of population contraction (driven by the temporary interruption and long term reduction of transmission of GR worms) outpace the rate at which the SOR population stabilises and expands (because they continue to be transmitted). If GR decline is faster than SOR expansion and the population threshold for maintenance of transmission is reached, local elimination will occur. If expansion of the SOR population prevents that threshold from being reached, transmission will persist, prevalence of infection will rebound and elimination will not occur. The outcome of the race will be determined by: (i) the starting frequency of SOR in a naïve population when ivermectin treatment begins, (ii) the relative rates of GR population contraction and SOR population expansion, and (iii) the population size at which transmission is no longer sustainable (the threshold for maintenance of transmission). Developing genotyping assays that can measure the pre-treatment (or current) SOR frequency and monitor the relative rates of GR contraction and SOR stabilization/expansion (the two critical parameters identified above) in populations undergoing CDTI is therefore essential to detect populations in which elimination may prove problematic. Furthermore, the response phenotypes in terms of dynamics of microfilariae production [36] and thus availability of skin microfilariae for vectors to ingest need to be better understood. They could then be taken into account for timing CDTI relative to vectors abundance so that the ratio of progeny of SOR/progeny of GR is minimal when the availability of vectors is maximal.

The third crucial question is whether SOR is likely to spread from one location to another. The analysis of population structure suggests that all of the communities sampled in Ghana are in a single transmission zone (there is no internal population structure between the Ghanaian communities), so that SOR could be transmitted between communities in this region of Ghana. This is also likely true for Cameroon. The tentative conclusion from analysis of the Sequenom data for MBM94, MBM07 and NKA07 from Cameroon suggests that immigration from an area not under CDTI (Nkam Valley) into a population under CDTI may occur (suggested by the similarity between MBM07 and NKA07), even if the pre-treatment level of transmission between the two populations is low (as is suggested by comparison of NKA07 and MBM94, which are genetically distinct populations). While the concept of recrudescence in areas undergoing control as a result of immigration from neighbouring regions with less or no control is not new, these data provide the first genetic evidence that such immigration has likely occurred and may have had an impact on the success of ivermectin distribution in the Mbam valley parasite populations. In contrast, Ghana and Cameroon are clearly separated genetically, which implies there is no transmission of *O. volvulus* at this scale and SOR will not spread from Ghana to Cameroon (or *vice versa*). While this is not surprising given the location of the two countries, the data show that genetic markers were able to detect transmission between endemic areas within Cameroon (i.e. are transmission zone markers) and thus could be a useful tool for onchocerciasis control programs. Such a tool could support decisions on whether to stop CDTI when criteria for stopping CDTI have been met in one area but not in others in a geographic context that makes transmission between these two areas possible. These data also suggest that not only can genetic markers sensitively detect transmission between geographic locations, but also that QLTs that are associated with SOR could increase in frequency in neighbouring regions having little drug selection pressure as a result of that transmission.

With respect to SORs, genotyping assays will benefit control programs in multiple ways by (i) providing diagnostic tools to monitor changes in the frequency of SORs (e.g. genotyping infective larvae in the vector), (ii) discriminating between genetic explanations for persistence of transmission (i.e., selection for SOR) and other factors that determine CDTI success (such as host-related factors, treatment coverage and compliance, pre-control prevalence and intensity of infection and vector biting rates [48–53, 99]), and (iii) suggesting a trigger for the initiation of alternative treatment strategies, such as anti-Wolbachia treatment [101, 102], local vector control, or new treatments that may become available sooner or later like moxidectin [46, 103], or combination treatments, flubendazole or emodepside (as reviewed in [104]) in populations where persistent transmission is observed in spite of CDTI [105].

### Summary

The data presented suggest that the evolution of SOR to ivermectin in *O. volvulus* is via soft selective sweeps of pre-existing quantitative trait loci (QTLs) rather than via a hard selective sweep of a relatively rare resistance-conferring mutation. The outcome of this soft selection is the accumulation of many alleles in a limited number of functional pathways that facilitate the recovery of adult female worm fecundity from the inhibitory effects of ivermectin. This is consistent with the observation that the acute microfilaricidal and macrofilarial anti-fecundity effects of ivermectin remain unaltered in SOR populations, and that the difference between SOR and GR parasites is quantitative variation in the rate and extent to which fertility recovers after ivermectin treatment. This conclusion (of a soft selective sweep of quantitative variation in the rate of recovery from ivermectin toxicity) is based on the presence of multiple, geographically independent genetic signals throughout the genome that differentiate GR and SOR pools and the apparent preservation of genetic diversity within the SOR populations following selection. Furthermore, when these data are considered together with the population bottlenecks that characterise the transmission of *O. volvulus* through its lifecycle and the likelihood of some degree of inbreeding, we conclude that *O. volvulus* populations are variable and can be structured due to allele frequency change in the absence of selection, i.e., as a consequence of genetic drift. What is unclear is the extent to which each putative QTL contributes to the development of SOR. The failure to detect differences between SOR and GR worms using a panel of 130 SNPs does suggest that the signature of selection in these populations is subtle and that no single putative QTL dominates the response; further validation of the putative QTLs is clearly required. A more thorough association analysis of each genomic region with SOR may be achieved by additional whole genome sequencing using single adult worms (as opposed to pool sequencing as described here) together with a more precise estimation of the ivermectin response of the individual worm. This would allow fine mapping of QTLs, and estimation of their effect size and penetrance, and ultimately, of the heritability of SOR. Such analyses should take transmission zones and population stratification into account to correctly determine the extent of gene flow and therefore, the ability of SOR alleles to be transmitted within and between populations. Additional sequencing of individual parasites may also provide greater confidence in assigning genetic associations to individual SNPs, opening the way to investigation of the likely function of individual SNPs as putative SOR-conferring variants. Given the increasing accessibility to genomic resources, the reduced cost of next-generation sequencing, and the ability to look at the whole genome in an unbiased way, we propose that genome-wide analyses such as those applied here should replace candidate gene approaches for future work concerned with the genetics and diagnosis of drug resistance in helminth parasites.

## ABBREVIATIONS

CDTI: community-directed treatment with ivermectin
GR: good responders
HWE: Hardy-Weinberg equilibrium
KS: Kolmogorov-Smirnov
MDS: multidimensional scaling
NLT: naïve or little treated
QT: quantitative trait
QTL: quantitative trait loci
SD: standard deviation
SNP: single nucleotide polymorphism
SOR: sub-optimal responders

## ACKNOWLEDGEMENTS

We would like to thank all of the patients and collaborators from Cameroon and Ghana who participated in the studies, as well as the Ministry of Public Health from Cameroon and the Ministry of Health from Ghana. We would like to acknowledge Pierre Lepage, Rosalie Frechette, Audrey Ann Kustec, Patrice Charbonneau-Larose and Valérie Catudal from the Genome Quebec Innovation Centre, Andrew Robinson and Nathan Hall from La Trobe University for computational support, James Cotton and Shannon Hedtke for critical reading of the manuscript, and Matthew Berriman and James Cotton from the Wellcome Trust Sanger Institute for pre-publication access to *O. volvulus* genomic resources.

## AUTHOR CONTRIBUTIONS

Conceived and designed the experiments: WNG, RKP, MB, CB Conducted the fieldwork, collected and processed parasite material for genetic analysis: MB, MYOA, SW, SDSP, CB, HCND, JAKO, JB, HC Contributed reagents and materials: MYOA, SW, MB, SDSP, RKP Led the parasitology and molecular biology: CB Led the bioinformatics and population genetics analyses: SRD Provided intellectual input, participated in meetings and discussions on the interpretation and presentation of the results and contributed to the writing of the manuscript: SRD, CB, HCND, JAKO, SDSP, JK, SW, ACK, MW, MGB, DAB, MYOA, MB, RKP, WNG Drafted the paper: SRD, WNG Read and approved the final submitted manuscript: SRD, CB, HND, JAKO, SDSP, JB, JK, SW, HC, ACK, MW, MGB, DAB, MYOA, MB, RKP, WNG

## SUPPLEMENTARY DATA

**Additional file 1 (xls).**

**Table S1.** Sequencing library composition, sequencing data and mapping statistics.

**Table S2.** SNP calling statistics derived from CRISP, FreeBayes and PoPoolation2 Pool-seq variant analysis, highlighting total SNPs called and the intersection between different SNP callers.

**Table S3.** Analysis of shared regions of genetic differentiation between SOR and GR of both Cameroon and Ghana. Table provides details of variants presented in Figure 1 A and B.

**Table S4**. Characterisation of QTL clusters that differentiate SOR and GR in either Cameroon or Ghana. Table provides details of regions of differentiation presented in Figure 1 C&D.

**Table S5.** Characterisation of genes within QTL clusters that differentiate SOR and GR in either Cameroon or Ghana.

**Table S6**. Analysis of genetic differentiation between GR and SOR pools in ivermectin-resistance associated genes from the literature and in clusters.

**Table S7**. Names and coordinates of the 160 SNPs genotyped by Sequenom.

**Table S8**. HWE analysis of GR and SOR populations from Cameroon.

**Table S9**. HWE analysis of GR and SOR populations from Ghana

**Additional file 2**: Supplementary information (pdf)

## Section 1. Extended information regarding sample history

**Table S1.** Criteria for phenotypic classification of *Onchocerca volvulus*

**Table S2.** Overview of samples selected for Pool-seq

**Table S3.** Overview of samples selected for single worm genotyping

**Figure S1.** Maps of sampling sites in Ghana and Cameroon

**Table S4.** Sampling sites and mapping coordinates in Ghana and Cameroon.

## Section 2. Extended discussion of hard-and soft-selective sweeps

**Figure S2.** Demonstrating the consequences of hard-versus soft-selection on genetic diversity

**Table S5**. A simple multilocus quantitative trait model demonstrating multiple genotypes conferring the same quantitative phenotype

**Table S6.** Summary of features differentiating hard and soft selection

## Section 3. Population assignment based on Sequenom genotyping of individual worms

**Table S7.** Population assignment of individual worms based on their genotype profile from 130 SNPs analysed by Sequenom

**Figure S3.** DAPC analysis of genetic diversity to compare predicted versus known population assignment of individuals based on their genotype.

